# Bioacoustic Event Detection with Self-Supervised Contrastive Learning

**DOI:** 10.1101/2022.10.12.511740

**Authors:** Peter C. Bermant, Leandra Brickson, Alexander J. Titus

## Abstract

While deep learning has revolutionized ecological data analysis, existing strategies often rely on supervised learning, which is subject to limitations on real-world applicability. In this paper, we apply self-supervised deep learning methods to bioacoustic data to enable unsupervised detection of bioacoustic event boundaries. We propose a convolutional deep neural network that operates on the raw waveform directly and is trained in accordance with the Noise Contrastive Estimation principle, which enables the system to detect spectral changes in the input acoustic stream. The model learns a representation of the input audio sampled at low frequency that encodes information regarding dissimilarity between sequential acoustic windows. During inference, we use a peak finding algorithm to search for regions of high dissimilarity in order to identify temporal boundaries of bioacoustic events. We report results using these techniques to detect sperm whale (*Physeter macrocephalus*) coda clicks in real-world recordings, and we demonstrate the viability of analyzing the vocalizations of other species (e.g. Bengalese finch syllable segmentation) in addition to other data modalities (e.g. animal behavioral dynamics, embryo development and tracking). We find that the self-supervised deep representation learning-based technique outperforms established threshold-based baseline methods without requiring manual annotation of acoustic datasets. Quantitatively, our approach yields a maximal R-value and F1-score of 0.887 and 0.876, respectively, and an area under the Precision-Recall curve (PR-AUC) of 0.917, while a baseline threshold detector acting on signal energy amplitude returns a maximal R-value and F1-score of 0.620 and 0.576, respectively, and a PR-AUC of 0.571. We also compare with a threshold detector using preprocessed (e.g. denoised) acoustic input. The findings of this paper establish the validity of unsupervised bioacoustic event detection using deep neural networks and self-supervised contrastive learning as an effective alternative to conventional techniques that leverage supervised methods for signal presence indication. Providing a means for highly accurate unsupervised detection, this paper serves as an important step towards developing a fully automated system for real-time acoustic monitoring of bioacoustic signals in real-world acoustic data. All code and data used in this study are available online.

## 1 Introduction

As conservation and management strategies encompass progressively more extreme solutions, ranging from species de-extinction to translation of non-human communication^1^, novel computational techniques can provide improved methods for processing large quantities of multimodal ecological data. In recent years, advances in machine learning (ML) and deep learning (DL), in particular, have revolutionized the analysis of bioacoustic data, facilitating the development of automated pipelines for processing data and contributing to enhanced conservation tactics across diverse taxa. To date, applications of DL to bioacoustics tend to rely on heavily supervised methods demanding large manually labeled datasets^2^, and these approaches often treat detection as a binary classification task^3^ that is more reflective of presence indication than event detection^2^. In this paper, we propose a self-supervised representation learning-based approach to bioacoustic event detection and segmentation that can predict temporal onsets and offsets of bioacoustic signals in a fully unsupervised regime.

Recent developments in observation hardware and methods now offer ecologists and biologists immense quantities of data captured via high-end cameras, static microphones, biologging devices, and satellites, among others^2^. The unprecedented amounts of high-quality data supersede the conventional approach of manual annotation and demand the application of novel computational methods to discover patterns of animal behavior with important ecological implications. To this end, contemporary methods of data analysis–and bioacoustic data analysis, in particular–often leverage DL and deep neural networks (DNNs).

In general, implementations of ML to ecological data analysis heavily depend on supervised techniques in which sufficiently large training datasets have been curated with accurate information, often corresponding to labels such as presence, location, or other identifying features of a species, object, sound, etc. For example, early research studies applied DL to sperm whale bioacoustics^4^, using supervised learning to train DNNs to perform detection and classification tasks of input spectrogram feature representations, requiring manually labeled datasets annotated according to relevant targets including but not limited to signal presence, clan membership, individual identity, and call type. Large-scale studies often employ Convolutional Neural Network (CNN) architectures to carry out supervised detection or classification tasks across diverse species. These include detection of humpback whale vocalizations in passive acoustic monitoring datasets^5^; detection, classification, and censusing of blue whale sounds^6^; avian species monitoring based on a CNN trained to predict the species class label given input audio^7^; presence indication of Hainanese gibbons using a ResNet-based CNN^8^; and, recently, detection and classification of marine sound sources using an image-based approach to spectrogram classification^9^. However, such supervised learning approaches remain limited in their scope, hindering their capacity to be deployed in real-time data processing pipelines. A salient limitation remains the dependency on large amounts of high-quality manually annotated datasets^2^, a problem compounded by its reliance on trained experts, the laborious nature (i.e. extensive time and resources) involved in manual labeling, and the potential for bias, variability, or uncertainty during the labeling process associated with variable human confidence and perception, especially across cohorts of individual annotators^10,11^. Further, manual data annotation remains expensive in terms of time^2^, expertise, and financial cost^12^, which has motivated citizen science-based crowdsourcing efforts to annotate large datasets at scale, though these approaches are similarly accompanied by error and limitations^13,14^. Reliance on manual annotations represents a key constraint on supervised DL methods for bioacoustic data analysis, making unsupervised approaches an attractive alternative with the potential to provide greater insight into animal communication and behavior.

More fundamentally, however, the foundational principles upon which DNN classifier-based detectors are constructed limits their applicability to real-world ecological data analysis. DNN-based bioacoustic detectors often pose the detection problem as a binary classification problem^5^ in which a neural network learns discriminative feature representations of input acoustic features (e.g. raw waveforms, handcrafted acoustic parameters, spectrograms, miscellaneous time-frequency representations), enabling the model to predict a binary class label denoting signal or non-signal (i.e. background). While attempts to address this shortcoming exist, often by employing smaller window (i.e. acoustic segment) sizes to artificially improve temporal resolution^15^, these approaches continue to treat detection as a presence indication task, unable to localize the signal directly within the window. Further, this workaround method of smaller windows is accompanied by its own disadvantages, mainly an impaired computational efficiency due to the need for overlapping windows to account for signals that could occur at the transition boundary between adjacent spectrograms^2^. The inability to precisely locate signal in a continuous recording renders it challenging to address downstream questions of ecological and/or animal behavioral importance. This is especially true in unsupervised regimes when labels (individual identity, call type, etc.) may not be readily accessible. For instance, unsupervised clustering-based techniques for call-type classification^16^ often expect manually segmented calls, which remain a challenge to extract from DNN-based presence indicators.

As an alternative approach, we treat the detection problem as a segmentation problem in which we aim to predict temporal onsets and/or offsets of bioacoustic event signals given a continuous stream of acoustic data. While previous studies have attempted both supervised^17–19^ and unsupervised^20,21^ bioacoustic sound event detection (SED) and segmentation, modern DNN-based–and self-supervised–methods remain underexplored^3^. We take inspiration from recent advances in representation learning and self-supervised contrastive learning^22–25^, especially those applied to human phoneme boundary detection and segmentation^26,27^. We propose a fully unsupervised DNN-based system for bioacoustic signal event detection that leverages self-supervised contrastive learning to map input audio to a learnable feature representation that encodes information regarding spectral boundaries, transitions, and changes that distinguish non-signal background from signal events.

Finally, we adjust our methods to account for unique challenges incurred by processing real-world bioacoustic data recorded in the native environment. In particular, a key challenge remains the presence of significant background noise in real-world bioacoustic recordings and the associated difficulties DNNs have in processing noisy data–often requiring modifications to DL-based methods to accommodate non-zero noise^28^–and distinguishing signal from non-signal^29^, especially in unsupervised schemes. As in previous work^30^, we account for environmental noise that would otherwise interfere with the contrastive learning objective by integrating on-the-fly noise reduction layers into the model architecture. In this paper, we provide an unsupervised framework for bioacoustic event detection based on contrastive representation learning, and we design our methods to operate directly on real-world acoustic data.

## 2 Materials and Methods

Motivated by recent advances in acoustic and visual representation learning involving self-supervised contrastive training objectives, we apply representation learning techniques to bioacoustic data to enable unsupervised detection and segmentation of bioacoustic events. In particular, given a continuous stream of unlabeled acoustic data, we implement a self-supervised learning algorithm that exploits an auxiliary proxy task with pseudo labels inferred from unlabeled data^25^ and aims to train a model, *f*, to encode the raw input waveform to a spectral representation that emphasizes spectral boundaries in the signal. The method relies on the Noise Contrastive Estimation principle^31^, which involves optimizing a probabalistic contrastive loss function that allows the model, *f*, to distinguish between sequential acoustic windows and randomly sampled distractor windows. For the model, *f*, we make use of a deep neural network to discover hidden patterns and complex relationships^32^ and to extract features relevant for optimizing the contrastive learning objective. During inference, a peak-finding algorithm detects regions of high dissimilarity between the encoded features of adjacent acoustic windows, which correspond to temporal boundaries of bioacoustic events.

### 2.1 Dataset

We apply our methods to a sperm whale (*Physeter macrocephalus*) click dataset. We process the raw acoustic data using the ‘Best Of’ cuts from the William A. Watkins Collection of Marine Mammal Sound Recordings database from Woods Hole Oceanographic Institution (https://cis.whoi.edu/science/B/whalesounds/index.cfm), retrieving 71 wav files amounting to 1.5 hours of acoustic data. Of these, we manually label with high confidence the clicks present in 38 files using Raven Pro 1.6.1, yielding a dataset containing 1738 annotated transients. All audio files were resampled to 48 kHz.

### 2.2 Self-Supervised Feature Extraction

Operating on the raw acoustic waveform directly, the model maps the input audio to a representation encoding discriminative features between sequential acoustic windows. Feature extraction involves training a neural network, 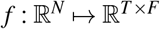, that encodes the *N*-sample acoustic waveform to a *T*-element temporal sequence of *F*-dimensional spectral features. While the model can process variable-length input waveforms, during training, the model *f* is given a real-valued continuous vector with fixed length *N*, i.e. **x** ∈ ℝ^*N*^, that represents the input audio signal and outputs an encoded spectral representation **z** = (**z**_1_,…,**z**_*T*_) ∈ ℝ^*T*×*F*^ consisting of a sequence of feature vectors sampled at low frequency. To train *f*, we follow the paradigm of Kreuk et al., 2020^26^. We optimize the encoding function *f* in accordance with the Noise Contrastive Estimation principle such that the network learns to maximize the similarity between adjacent (i.e. sequential) windows **z**_*i*_, **z**_*i*+1_ for *i* ∈ [1, *T*] in the learned representation while minimizing the similarity between randomly sampled distractor windows **z**_*i*_, **z**_*j*_ for **z**_*j*_ ∈ *D*(**z**_*i*_), the set of nonadjacent windows to **z**_*i*_, i.e. *D*(**z**_*i*_) = {**z**_*j*_ : |*i* – *j*| > 1, **z**_*j*_ ∈ **z**}.

Following Kreuk et al., 2020^26^, we use the cosine similarity between two feature vectors as our similarity metric

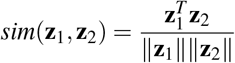

For the contrastive loss function, we implement a cross-entropy loss

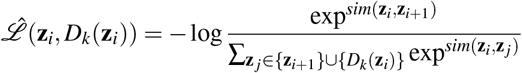

This means that given a batch of *n* training samples 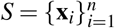, the total loss is given by^26^

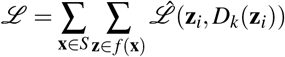

Given an acoustic waveform as input, this training criterion seeks to yield a representation in which the dissimilarity between sequential encoded feature vectors of (i.e. adjacent) audio windows is minimized.

### 2.3 The Model

We use a Convolutional Neural Network (CNN) comprised of one-dimensional convolutional layers (Conv1d) that operate on the raw waveform directly. Importantly, we also include on-the-fly data preprocessing layers to remove acoustic interferences.

In particular, the first layer of the model *f* consists of a high-pass filter layer constructed using a nontrainable Conv1d layer with frozen weights determined by a windowed *sinc* function^30,33,34^. Given the broadband nature of sperm whale clicks^35^, we select a cutoff frequency of 3 kHZ and a transition bandwidth 0.08 to remove low-frequency environmental background noise while preserving the spectral structure of the clicks.

Following the denoising layer is the convolutional model, consisting of Conv1d layers with batch normalization (Batch-Norm1d) and LeakyReLU activation. We use a 4-layer network with kernel sizes [8, 6, 4, 4] and strides [4, 3, 2, 2], corresponding to a hop length of 48 samples (i.e. 1ms) and a receptive field of 136 samples (i.e. ~3ms). For all Conv1d layers, we use 128 output channels. Lastly, we include a fully-connected linear projection layer with Tanh activation to output the spectral representation. With this architecture, the model *f* encodes input audio to a learned representation comprised of a sequence of 32-dimensional feature vectors sampled at 1 kHz.

To train the model, we optimize the contrastive learning objective using Stochastic Gradient Descent (SGD). We use a learning rate 1e-4, momentum 0.9, batch size 64 for 50 epochs. Given the stochasticity of training, we serialize the model weights after each epoch and select the top-performing (in terms of the learning objective) model for inference.

The baseline energy amplitude detector model (which is functionally equivalent to a signal-to-noise (SNR)-based threshold detector^36^) requires no trainable parameters and involves using a peak-finding algorithm to detect peaks in the energy of the input waveform. We use the raw audio as input. We repeat the baseline detector with and without data preprocessing (e.g. high pass filter denoising). In this case, all suprathreshold detections are considered to be clicks.

### 2.4 Inference and Peak-Finding

During inference, we obtain the spectral representation **z** given the acoustic waveform **x** using the trained model (i.e. **z** = *f* (**x**)). We compute the distance between sequential windows using a bounded L2 dissimilarity metric:

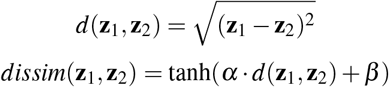

where *α* and *β* are fixed hyperparameters obtained by exhaustive grid search. Unlike the cosine similarity metric, which neglects normalization, the Euclidean distance-based metric preserves information regarding scaling, which is an important consideration for signals in the presence of low-amplitude nonstationary noise. We evaluate the dissimilarity between the feature representations of adjacent windows *dissim*(**z**_*i*_, **z**_*i*+1_) for *i* ∈ [1, *T* – 1]. As the contrastive learning objective aims to suppress dissimilarity between successive windows, a relatively high dissimilarity metric indicates a spectral boundary, suggesting a transition from background to bioacoustic signal (or vice versa) in the input audio. In the case of sperm whale clicks, we smooth the dissimilarity metric over the temporal axis by convolving *dissim* with a box function of width *σ*, operating under the assumption that we are interested in single click times as opposed to boundaries corresponding to onsets and offsets of the transient clicks.

Finally, using the `scikit-learn` package, we employ a peak-finding algorithm to search for peaks in the dissimilarity. We search over peak prominences *δ* and attribute all suprathreshold peak detections to sperm whale clicks.

### 2.5 Evaluation Metrics

Motivated by work on phoneme boundary detection^26^, we evaluate model performance using precision (P), recall (R) and F1-score with a tolerance level *τ*. As in Kreuk et al., 2020^26^, we also include the R-value, which serves as a more robust metric than F1-score that is less sensitive to oversegmentation. In the case of sperm whale clicks we choose τ = 20 ms, a value that exceeds observed inter-pulse intervals (IPIs)^37^ but is less than typical inter-click intervals (ICIs)^38^, even in the case of buzzes which can exhibit click rates 1-2 orders of magnitude larger^39^. This emphasizes the objective of resolving acoustic structures on the temporal order of individual clicks. Finally, we report the area under the Precision-Recall curve (PR-AUC).

## 3 Results

In Table 1, we report F1-score, R-value, and PR-AUC for the proposed self-supervised detection model as well as the energy threshold detector baseline with and without high pass filter (HPF) preprocessing. Our method achieves a maximal F1-score of 0.876, an R-value of 0.887, and a PR-AUC of 0.917. The baseline threshold-based energy detection method operating on raw input audio with no data preprocessing (e.g. denoising) yields an F1-score of 0.576, an R-value of 0.620, and a PR-AUC of 0.571. Inclusion of denoising in the threshold-based energy detection method yields modest performance benefits by eliminating high-amplitude low frequency environmental noise, boosting the F1-score, R-value, and PR-AUC to 0.639, 0.643, and 0.706, respectively. These results suggest that the unsupervised deep learning-based approaches to bioacoustic signal detection implemented in this paper can outperform established baseline techniques for detecting events in acoustic recordings. PR curves are presented in Fig. 3.

**Table 1.** Summary of experiments implemented in this study. We compare our approach based on self-supervised learning (SSL) with baseline energy amplitude threshold detectors with and without high pass filter (HPF) preprocessing. We report PR-AUC, maximal F1-Score, and maximal R-value.

| Model | PR-AUC | F1-Score | R-Value |
| --- | --- | --- | --- |
| SSL Detection | 0.917 | 0.876 | 0.887 |
| Energy Amplitude Detection | 0.571 | 0.576 | 0.620 |
| Energy Amplitude + HPF Detection | 0.706 | 0.639 | 0.643 |

In Fig. 1, we demonstrate the self-supervised detection pipeline and results applied to an acoustic recording containing a sequence of clicks. Fig 1 **a**) represents the acoustic waveform, on which the model *f* operates directly. Fig 1 **b**) presents a spectrogram representation to visualize the spectrotemporal features of the signal. The learned representation **z** resulting from the optimization of the contrastive learning objective is shown in Fig 1 **c**). Finally, we compute the bounded L2 dissimilarity metric using *α* = 0.1 and *β* = 0.9 and plot the result in Fig 1 **d**). Employing a peak-finding algorithm, we search over prominences *δ* to obtain the PR-AUC, observing maximal F1-score with *δ* = 0.1, and we superimpose the detected peaks in Fig 1 **d**). Together, the quantitative metrics and visualizations in Fig 1 demonstrate the efficacy of applying self-supervised contrastive learning in the context of bioacoustics event detection.

**Figure 1.**
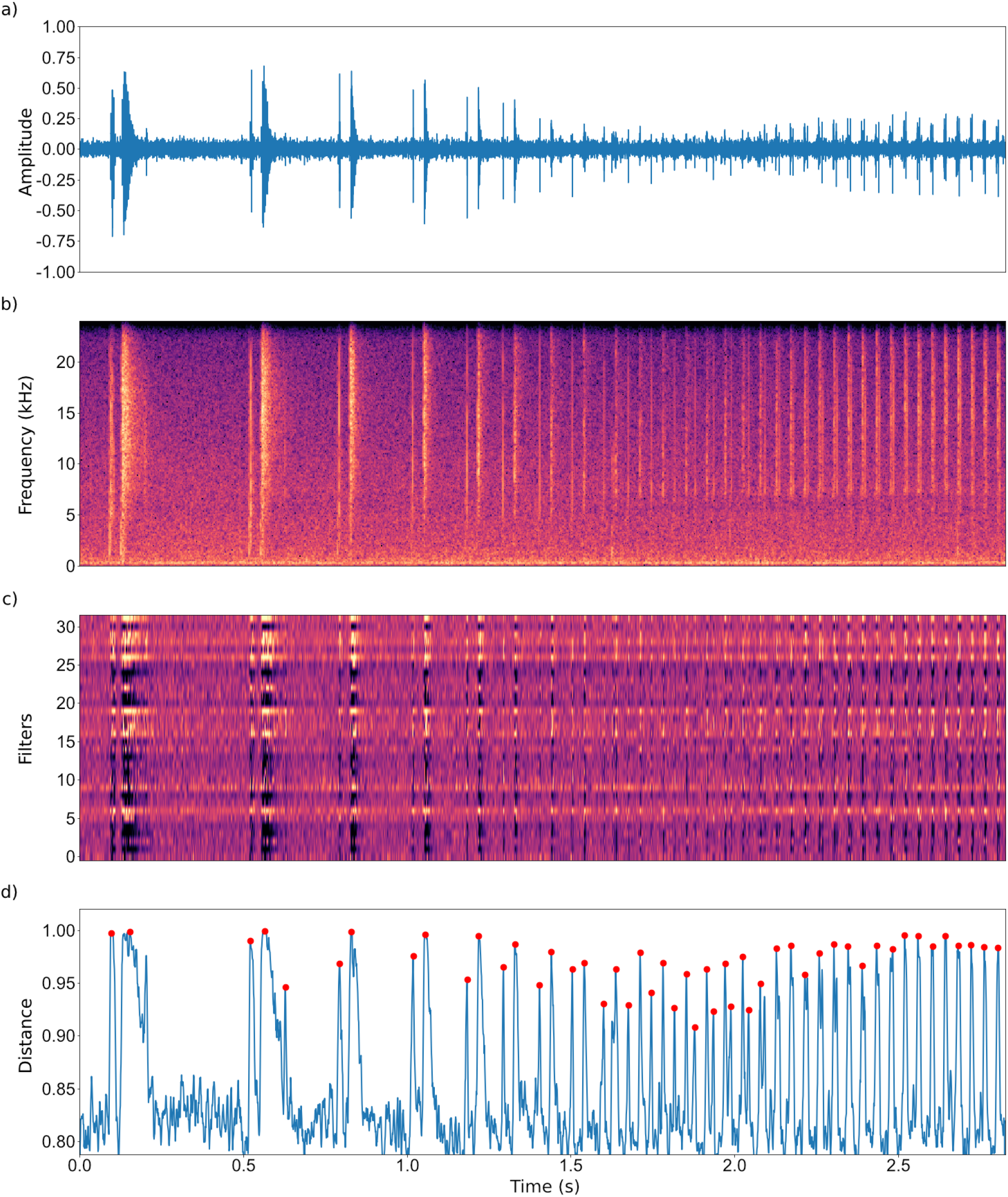
Application of self-supervised contrastive learning to sperm whale click detection. (**a**) Input acoustic waveform. (**b**) Spectrogram representation. (**c**) The learned representation. (**d**) The distance metric *dissim* demonstrating boundaries of high dissimilarity between acoustic windows. Peaks were detected with α = 0.1, *β* = 0.9, σ = 5ms, δ = 0.1

In Fig. 2, we provide initial exploratory results applying self-supervised contrastive learning to other datasets and data modalities. In Fig. 2 **a**), we segment Bengalese finch (*Lonchura striata domestica)* birdsong^40^ into predictions for sequential vocal units, or syllables. In Fig. 2 **b**), we use the contrastive detection framework to predict behavioral transitions in green sea turtle (*Chelonia mydas*) behavioral dynamic data^41^. In Fig. 2 **c**), we investigate unsupervised embryo tracking by identifying transitions characteristic of cell divisions in a mouse embryo development dataset^42^. We defer a more thorough analysis to future studies.

**Figure 2.**
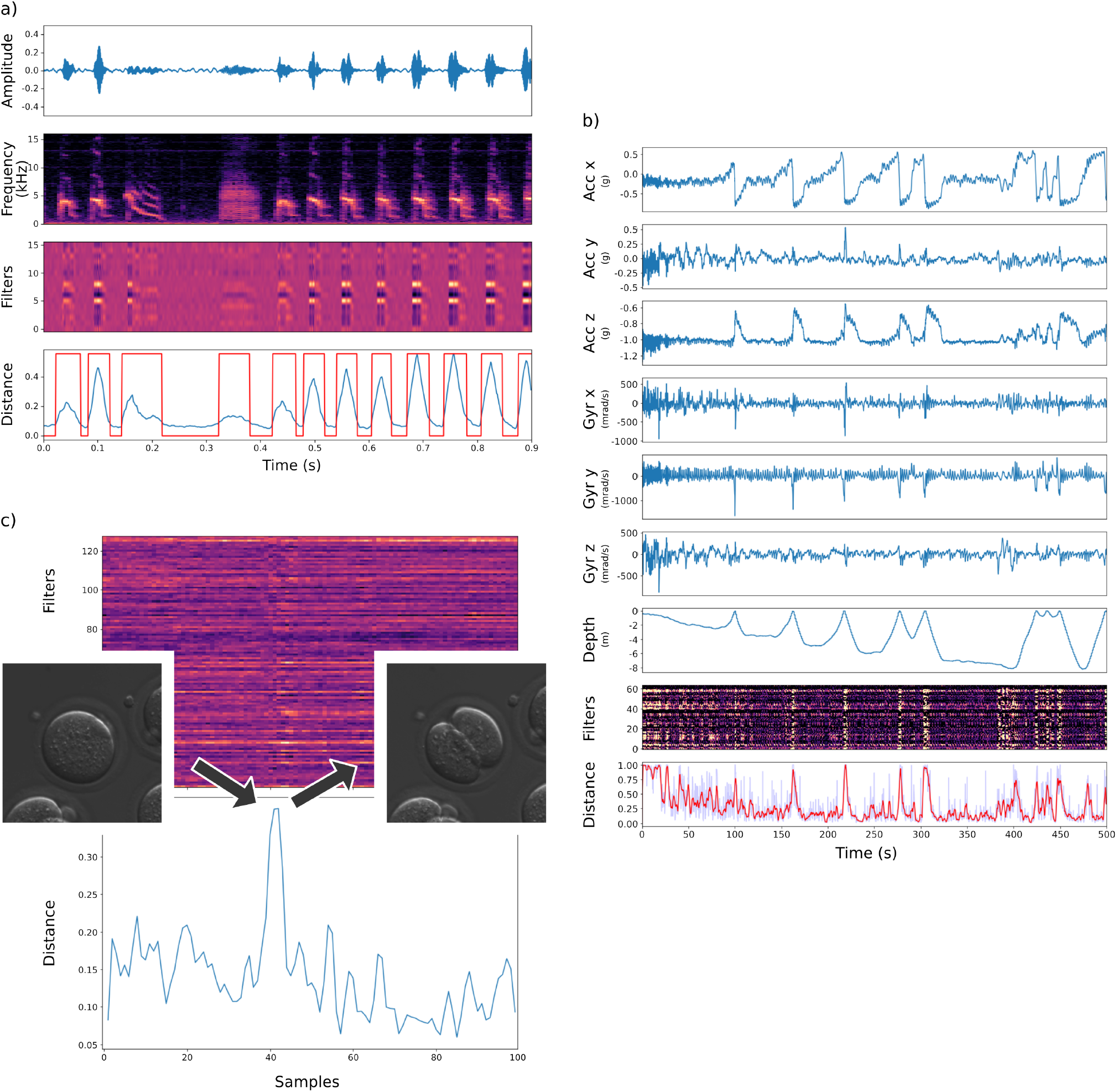
Schematic experimental applications of self-supervised contrastive learning to other datasets and data modalities. (**a**) Bengalese finch syllable segmentation; vocal units were segmented using a Z-score-based peak detection algorithm; (**b**) Detection of behavioral transitions in free-ranging Green sea turtles according to animal-borne multi-channel accelerometry and gyroscope data. (**c**) Tracking of mouse embryo development.

**Figure 3.**
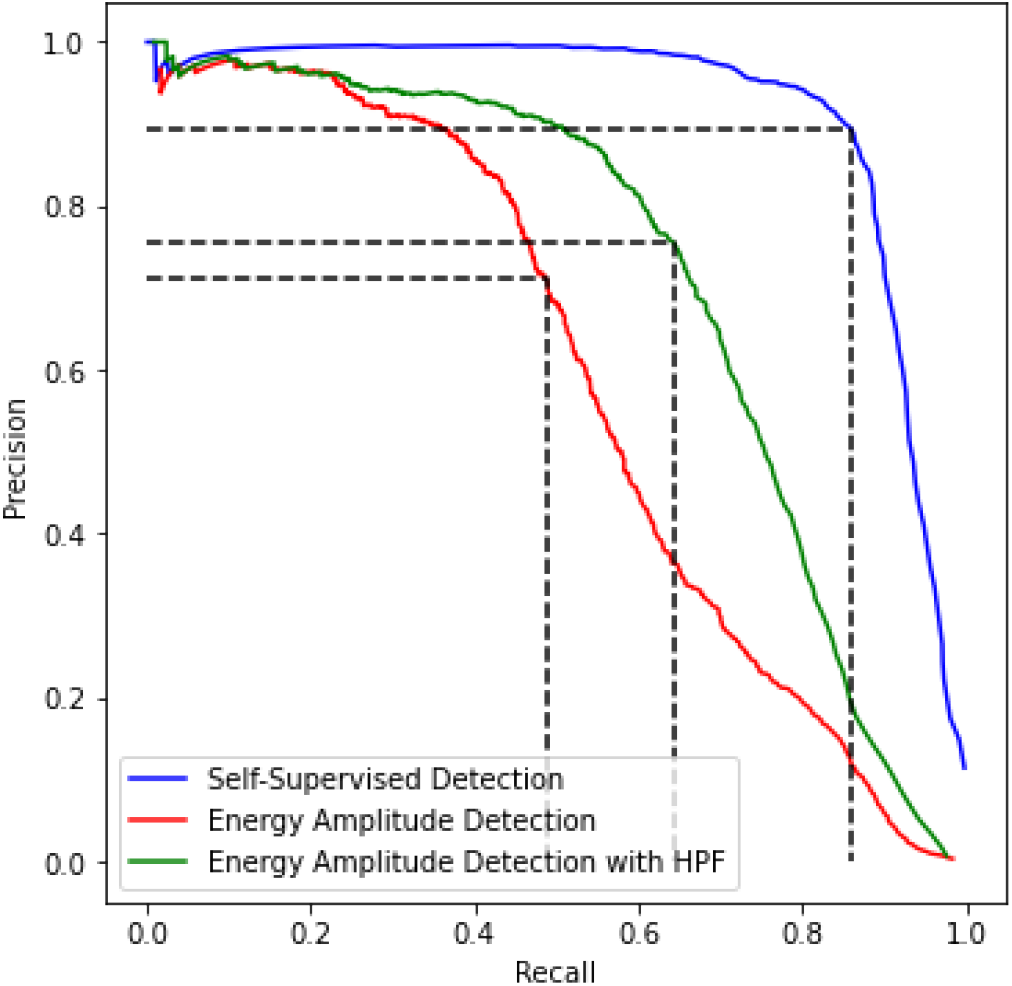
Precision-Recall (PR) curves for our self-supervised detector (blue), the baseline energy amplitude threshold detector with high pass filter (HPF) preprocessing (green), and the baseline energy amplitude threshold detector with no preprocessing (red). Dashed lines denote PR values corresponding to maximal F1-scores.

## 4 Discussion

While state-of-the-art computational methods for bioacoustic event detection tend to treat detection as a binary classification and/or presence indication problem and rely on supervised learning techniques, we propose a novel self-supervised approach to detect spectral changes associated with bioacoustic signals. Importantly, our unsupervised method requires no manual annotation during training, shifting the paradigm from existing techniques that expect large quantities of labeled training data. We demonstrate that unsupervised DNN-based approaches to bioacoustic detection and segmentation can outperform established baseline methods, yielding improved results in terms of precision, recall, F1-score, R-value, and PR-AUC.

### 4.1 Vision for a DL-Based Pipeline for Bioacoustic Monitoring

In line with existing research^21,29^, we envision a complete pipeline for ecological data analysis–particularly in the context of bioacoustics–involving discrete DNN-based modules to address ecologically important questions regarding animal communication and behavior. Further, we advocate the development and implementation of *un*supervised techniques to minimize the role of the human observer and circumvent the reliance on–and drawbacks of–manual annotation^2^. Explicitly, we propose a framework encompassing detection, classification, localization, and tracking (DCLT)^36^. The multi-step pipeline involves (1) signal processing; (2) event boundary detection; (3) source separation, localization, and tracking; and (4) clustering and classification. This study serves as a proof-of-concept to demonstrate the application of unsupervised DNN-based methods to solve Steps (1) and (2). We encourage future experimental setups to expand on our results.

Step (1) (i.e. signal processing) requires an emphasis on denoising given the infeasibility of obtaining non-noisy acoustic data in real-world environmental recordings; bioacoustic denoising is an active area of research^43^, and to our knowledge, there exist no DNN-based techniques for bioacoustic noise reduction, a regime that differs from the related subfield of human speech enhancement in that human speech enhancement typically relies on supervised methods to separate noise and signal^44^. While we integrate fixed denoising into our model architecture using a high-pass filter to eliminate low-frequency environmental noise, our methods could benefit from enhanced DL-based denoising to remove noise that spans the entire spectral zone of support for the given signal of interest.

The focus of our paper remains Step (2) (i.e. boundary detection). While our results indicate that DNN-based approaches can significantly outperform threshold-based methods (which were used in Koumura & Okanoya, 2016), more recent DL techniques could be employed to further boost performance. For instance, vector-quantized contrastive predictive coding (VQ-CPC) has been used to advance phoneme segmentation and acoustic unit discovery^27^.

Step (3) (i.e. bioacoustic source separation) has seen major inroads in recent years^30,45^. This includes the development of unsupervised source separation methods^45^, an important consideration that can improve downstream classification performance. While mixture invariant training (MixIT) algorithm^45^ provides an effective means for separating sources in bioacoustic recordings, bioacoustic source separation could further benefit from alternative unsupervised methods. The related problems of source localization and source tracking involve extracting signals from continuous acoustic streams, even when the sources responsible for generating the event are traveling in space and time; while DL-based approaches to localization have indicated promising results^3,46^, together, separation, localization, and tracking remain underexplored in the literature but represent important considerations when processing ecological data to assess animal behavior and communication^36^.

Step (4) (i.e. clustering and classification) can provide important insight into questions concerning animal behavior, communication, and ecology. Analyzing vocal repertoires, classifying vocalizations according to type, identifying unique individuals, and characterizing animal behavioral dynamics represent central classification-based objectives with significant ecological and conservation implications. While existing methods have explored both supervised^4^ and unsupervised^16^ approaches for classifying input data according to target label (i.e. call type, animal identity, etc.), DNN-based advances could offer improved performance and reduced reliance on the human observer. DNNs are powerful instruments for discovering hidden and cryptic patterns in large-scale multi-modal datasets^2^, and unsupervised DNN-based techniques for classification tasks, in particular, have shown the potential for enhancing clusterability and feature discrimination^47^, enabling the scientific community to answer complex questions regarding ecology and animal behavior.

### 4.2 Applications to Other Datasets and Data Modalities

Finally, in Fig. 2 we schematically demonstrate that these methods can generalize to other datasets and data modalities beyond bioacoustic click detection. In Fig. 2 **a**), we apply our methods to Bengalese finch^40^ syllable segmentation. Bengalese finch birdsong is comprised of sequential vocal elements, known as syllables. We reformulate the detection problem as a segmentation problem, which involves detecting temporal onsets and offsets of signals and extracting the vocal unit bounded by the detections. As in the case of sperm whales, we use a dataset containing acoustic recordings; in particular, we apply the contrastive learning objective to a collection of song from four Bengalese finches^40^ to segment the continuous audio stream into predictions for vocal units.

In Fig. 2 **b**), we use self-supervised learning to address green sea turtle behavioral dynamics^41^. The aim is to automatically identify, predict, and monitor various behaviors of free-ranging sea turtles in their natural habitat through the use of animal-borne multi-sensor recorders^48^. The dataset consists of multi-channel time series corresponding to acceleration, gyroscope, and depth recorded using animal-borne sensors from 13 immature green sea turtles. We show that self-supervised learning has the potential to reveal transition boundaries between behaviors, providing for automatic segmentation of animal behavior data. Future studies can consider downstream classification and/or clustering^16^ to predict the class label of the segmented behavior.

And in Fig. 2 **c**), we show that self-supervised contrastive learning can be leveraged to address questions regarding embryo tracking and development^42^. We use a mouse embryo tracking database containing 100 samples of embryos progressing to the 8-cell stage^42^. A DNN trained by optimizing the contrastive learning objective encodes embryo image input to a representation emphasizing boundaries and discontinuities in embryo development, which is used to predict transitions between developmental stages.

While we demonstrate the potential for additional implementations, we suggest further studies exploring ecological applications of our methods by employing self-supervised contrastive learning to detect discontinuities, transitions, and events in additional bioacoustics datasets as well as animal behavioral dynamics datasets.

## 5 Conclusion

In this paper, we apply self-supervised representation learning to ecological data analysis with an emphasis on bioacoustic event detection. In particular, we construct a CNN-based model and optimize a contrastive learning objective in accordance with the Noise Contrastive Estimation principle to yield a representation of input audio that encodes features involved in detecting spectral changes and/or boundaries, enabling the model to predict temporal onsets and/or offsets of bioacoustic signals. We compute a dissimilarity metric to compare sequential acoustic windows, and we employ a peak-finding algorithm to detect suprathreshold dissimilarities indicative of a transition from non-signal to signal and vice versa. In the case of sperm whale clicks, we present quantitative performance metrics in the form of F1-scores, R-values, and PR-AUCs, concluding that the unsupervised DL approach based on contrastive representation learning outperforms baseline methods such as energy amplitude detectors. Interestingly, we observe that omitting the smoothing operation σ may enable the resolution of finer-temporal-scale structures such as IPIs, which could allow biologists to identify individual sperm whales^49^ using an unsupervised computational technique; we encourage additional studies to further explore this observation. We also consider applications of the methods to other datasets and data modalities, including bioacoustic data produced by other taxa, behavioral dynamic data, and imaging data, demonstrating that the contrastive learning objective can have wide-ranging implications for ecological data analysis.

This paper serves as an important step towards the realization of a fully automated system for processing bioacoustic data while minimizing the conventional role of the trained expert human observer. Our methods and proposals pave the way for future studies that should aim to construct a complete framework for ecological data analysis in order to elucidate and understand animal behavior and, subsequently, to design better-informed strategies and approaches for species conservation at large.

## 6 Data Availability

The sperm whale click data that support the findings of this study are publicly available in the ‘Best Of’ cuts from the Watkins Marine Mammal Sound Database, Woods Hole Oceanographic Institution, and the New Bedford Whaling Museum (https://cis.whoi.edu/science/B/whalesounds/index.cfm). The bengalese finch data^40^ are available from *figshare* with identifier https://doi.org/10.6084/M9.figshare.4805749.V5, and the green sea turtle data^41^ used in this study are available from *Dryad* with identifier https://doi.org/10.5061/dryad.hhmgqnkd9. The embryo development data^42^ that support the results are publicly available online http://celltracking.bio.nyu.edu/. The custom-written code (Python 3.8.3) is available at our GitHub https://github.com/colossal-compsci/SSLUnsupDet.

## 7 Author Information

### Contributions

P.C.B performed task setting, data processing, machine learning, article writing, figure making, L.B. assisted in experimental design, A.J.T. supervised the analysis, and all authors wrote and reviewed the manuscript.

## 8 Competing Interests

The authors declare no competing interests.

## Notes

https://cis.whoi.edu/science/B/whalesounds/index.cfm

https://doi.org/10.6084/M9.figshare.4805749.V5

https://doi.org/10.5061/dryad.hhmgqnkd9

http://celltracking.bio.nyu.edu/

https://github.com/colossal-compsci/SSLUnsupDet

### Competing Interest Statement

The authors have declared no competing interest.

### Summary of Updates

Updating author affiliations

